# APOL1 variant-expressing endothelial cells exhibit autophagic dysfunction and mitochondrial stress

**DOI:** 10.1101/2020.03.18.996702

**Authors:** Ashira Blazer, Yingzhi Qian, Martin Paul Schlegel, Jill P. Buyon, Ken Cadwell, Michael Cammer, Sean P. Heffron, Feng-Xia Liang, Shilpi Mehta-Lee, Timothy Niewold, Sara E. Rasmussen, Robert M. Clancy

## Abstract

Apolipoprotein L1 (APOL1) gene risk variants (RV) associate with renal and cardiovascular disease particularly in SLE. We hypothesized that in RV-carrying human umbilical vein endothelial cells (HUVECs) cytokine-induced APOL1 expression compromises mitochondrial respiration, lysosome integrity, and autophagic flux. HUVEC cultures of each APOL1 genotype were generated. APOL1 was expressed using IFNɣ; HUVEC mitochondrial function, lysosome integrity, and autophagic flux were measured. IFNɣ increased APOL1 expression across all genotypes 20-fold (p=0.001). Compared to the homozygous G0 (ancestral) allele (0RV), high risk (2RV) HUVECs showed both depressed baseline and maximum mitochondrial oxygen consumption (p<0.01), and impaired mitochondrial networking on MitoTracker assays. These cells also demonstrated a contracted lysosome compartment (p<0.001), and an accumulation of autophagosomes suggesting a defect in autophagic flux. Treatment of 0RV HUVECs with a non-selective lysosome inhibitor, hydroxychloroquine, produced autophagosome accumulations similar to the 2RV cells, thus implicating lysosome dysfunction in blocking autophagy. Compared to 0RV and 2RV HUVECs, 1 RV cells demonstrated an intermediate autophagy defect which was exacerbated by IFNɣ. Our findings implicate dysfunction of mitochondrial respiration, lysosome, and autophagy in APOL1 RV-mediated endothelial cytotoxicity. IFNɣ amplified this phenotype even in variant heterozygous cells–a potential underpin of the APOL1/inflammation interaction. This is the first description of APOL1 pathobiology in variant heterozygous cell cultures.

## Introduction

Ancestrally African individuals, particularly those with autoimmunity, suffer from disproportionate rates of cardiovascular, hypertensive, and kidney disease. Two polymorphisms, G1 (SER342GLY; ILE384MET) and G2 (6BP deletion N388/Y389), of the Apolipoprotein L1 (APOL1) gene have been shown to associate with these adverse phenotypes in individuals of recent African heritage. These mutations have been evolutionarily conserved due to an advantage in resisting *Trypanosoma brucei*, the causal agent of African trypanosomiasis [1], and are therefore largely absent from non-African populations [2, 3]. It has been previously demonstrated that these variants are common in an African American lupus cohort with 53% of individuals heterozygous and 13% homozygous for the variants [4]. Despite the variants’ high allelic frequencies, the adverse phenotype penetrance varies considerably. It has been reported that endogenous and exogenous interferons, as seen in lupus, can precipitate the adverse phenotype in variant homozygotes [5].

Consistent with its innate immune function, APOL1 expression is highly responsive to inflammatory signals including Toll-like receptor (TLR) ligation and inflammatory cytokines such as tumor necrosis factor alpha (TNFα) and interferon gamma (IFNɣ) [6]. Immunoprecipitation assays show that interferon regulatory factor 1 and 2 and STAT2 bind the APOL1 promoter heightening expression [5, 7]. Therefore, APOL1 variant gene penetrance may be contingent upon environmental second hits [8]. Intracellularly accumulated APOL1 contains both a BH3 domain, which participates in initiating autophagy, and a pore-forming domain that can be inserted into phospholipid bilayers, causing tissue injury [9, 10]. This injury is contingent upon APOL1 protein accumulation beyond a toxic threshold. It may be of critical importance to consider that even heterozygous carriers express both ancestral and variant allele copies [11]. Current cell culture models introduce the variants using viral-vector systems hampering the ability to make inferences in the heterozygous state.

Several cell types including podocytes, human embryonic kidney cells, and oocytes over-expressing variant APOL1 have demonstrated mitochondrial injury, lysosome compromise, and autophagic flux defects resulting in cell death [12–16]. However, risk variant-mediated toxicity mechanisms have not been studied in vascular disease-relevant cell culture models [17]. Moreover, the G1 and G2 SNPs reside in amino acid coding regions therefore altering protein structural stability and function [18]. Despite this apparent gain of function property [19], the inheritance pattern is thought to be recessive, and the literature has not described variant-associated injury in heterozygous carrying tissues [20].

Endothelial dysfunction has been widely recognized as a risk factor for the development of vascular disease [21]. Accordingly, this study was initiated to address the hypothesis that in HUVECs, variant APOL1 confers mitochondrial stress, autophagy defects and loss of lysosome integrity--a phenotype heightened by exposure to an inflammatory milieu. Cytokine exposure may additionally drive APOL1 expression and amplify injury even in cells heterozygous for the variant. HUVECs were obtained from healthy controls of each genotype – homozygous ancestral allele (0RV), heterozygous (1RV), homozygous (2RV) – to determine the consequences of cytokine exposure across APOL1 genotype.

## Materials and Methods

### Human subjects

This study abides by the Declaration of Helsinki principles and was approved by the Institutional Review Board of New York University School of Medicine. Healthy pregnant women were recruited from a single center labor and delivery ward. Participants provided written informed consent for fetal umbilical cord collection. Inclusion criteria were: African ancestry (concordant partner), and age >18 years. Umbilical cords that could not be processed within two hours of delivery were excluded. In total, 15 cords were collected between February 2015 and December 2018. For experiments in which human sera were added to the HUVEC cultures (see below), samples were obtained from 5 SLE patients and 3 healthy controls. These subjects were enrolled between February 2015 and December 2018 in the NYU Division of Rheumatology-wide IRB-approved Specimen And Matched Phenotype Linked Evaluation (SAMPLE) biorepository. The SLE patients and healthy controls were African American and >18 years of age. Patients met at least 4 American College of Rheumatology criteria for SLE [22]. Clinical data at the time of sample draw included medications, ACR SLE criteria, autoantibody profile, and SLE disease activity score.

### HUVEC culture establishment and processing

Please see the Major Resources Table in the online-only Data Supplement for detailed descriptions of all antibodies and cultured cells used. A cut 5-cm section of umbilical cord was collected in RPMI media (Clonetics Corp.) supplemented with heparin 10 U/mL, penicillin/streptomycin 10 U/mL, and gentamycin 10 μg/mL. The umbilical vein was cannulated and perfused three times with HBSS solution to remove clotted blood. The umbilical vein was then perfused with collagenase A (type III) solution and both ends were clamped for 10 minutes to allow separation of umbilical vein cells as described [23]. Subsequently, the vein was re-cannulated and perfused again with HBSS, allowing cells to slowly drip from the vein into EGM-2 BulletKit Medium (Clonetics Corp.). The resulting solution was poured over a cell strainer. Cells were centrifuged and the pellet re-suspended in clean culture media (EGM-2 supplemented with 10% FBS, 50 U/ml penicillin, 100 mg/ml gentamicin). The cell isolate contained HUVECs, fibroblasts, and residual blood cells. To yield enriched cultures of HUVECs, the cell suspension was passed through a magnetic bead column to capture CD146+ cells. The residual filtrate was discarded. HUVEC cultures were expanded and passaged for use in these experiments described below. Using FACS analysis, HUVECs exhibited strongly positive staining for both CD31 and CD146. In total, 15 healthy HUVEC cultures were established representing genotypes as follows: 0RV n=8, 1RV n=4, 2RV n=3. There were no differences in donor infant gender distributions across genotype.

### APOL1 genotyping

To ensure cell cultures representing each genotype in triplicate were available for subsequent analysis, APOL1 genotyping was performed as described previously [4]. Briefly, genomic DNA was isolated from each HUVEC culture using the Qiagen kit (Valencia) according to the manufacturer’s instructions. DNA isolates were quantitated using a Nanodrop-1000 spectrophotometer (Nanodrop Products). One hundred ng of genomic DNA was used as a template for conventional polymerase chain reaction (PCR). A single 300 base-pair DNA segment containing the APOL1 polymorphisms, G1 (rs73885319 and rs60910145) and G2 (rs71785313), was amplified using AmpliTaq Gold 360 DNA Polymerase (Applied Biosystems). For quality control, DNA was elongated in both forward and reverse directions using sequences 5’-GCCAATCTCAGCTGAAAGCG-3’ and 5’-TGCCAGGCATATCTCTCCTGG-3’ respectively. Genotypes were analyzed using the GeneWiz online platform. Successful genotyping was completed on all DNA samples.

### Measurement of serum IFN-α activity

The reporter cell assay for IFN-α has been described in detail previously [24, 25]. In this assay, reporter cells are used to measure the ability of patient sera to cause IFN-induced gene expression. The reporter cells (WISH cells, ATCC #CCL-25) are cultured with 50% patient serum for 6 hours. The cells are lysed, and cDNA is made from total cellular mRNA and then quantified using real-time PCR. Forward and reverse primers for the genes IFN-induced protein with tetratricopeptide repeats 1 (IFIT1), myxovirus resistance 1 (MX1) and dsRNA-activated protein kinase (PKR), which are highly and specifically induced by IFN-α, were used in the reaction [24]. The relative expression of each of these three genes was calculated as a fold increase compared with its expression in WISH cells cultured with media alone and then standardized to healthy donors, and summed to generate a score reflecting the ability of sera to cause IFN-induced gene expression (serum IFNα activity) [24]. This assay has been highly informative in SLE and other autoimmune diseases [26–28].

### Inflammatory model of APOL1 expression

Single genotype HUVEC cultures grown in EGM-2 (Promocell, Heidelberg, Germany) supplemented with 10% FBS, 50 U/ml penicillin, and 100 mg/ml gentamicin were seeded at 20% confluence in 75 ml culture plates coated with 0.1% gelatin. Once confluent, cells were passaged and harvested once per week using trypsin-EDTA. Only HUVECs between passages 4-8 were used in experiments. For each genotype, untreated controls were compared to cells treated with one of the following: 50% human sera isolated from healthy donors or patients with SLE, IFNɣ (50pg/mL), IFNα (50pg/mL) or TNFα (50pg/mL). Cells were lysed; both protein and mRNA were extracted for immunoblot and qPCR.

### Immunoblot

Protein concentration was determined using a BCA protein assay kit (ThermoFisher Scientific) following manufacturer’s instructions. Appropriate concentrations of cell lysate were diluted with 4X Blot® LDS sample buffer then heated to 70°C for 5 minutes. Samples were resolved on Blot 4-12% Bis-Tris Plus Gels (Life Technologies) and transferred to PVD membranes. Membranes were blocked with Odyssey® Blocking Buffer (TBS) (Li-Cor Biotechnology) for 1 hour at room temperature. After blocking, membranes were incubated with rabbit anti-human APOL1 (1 μg/mL) (Sigma-Aldrich) and mouse anti-human tubulin (1 μg/mL) (AbCam) diluted in 5% BSA/TBS-T overnight. Membranes were then incubated with an HCRP-conjugated anti-mouse or anti-rabbit secondary antibody (1:2000) (Santa Cruz Biotechnology) for 1 hour at room temperature. Protein bands were visualized using Li-Cor Image Studio Lite 4.0. Immunoblots were quantified by densitometry of experimental bands relative to loading controls using ImageJ 1.51 Java 1.8 running on Windows 7 or 10.

### qPCR

Total RNA was extracted from endothelial cells using the RNAeasy Mini kit according to the manufacturer’s instructions (Qiagen). RNA was reverse transcribed to prepare cDNA libraries. Both forward and reverse primers for APOL1 were used (sequences above). Levels of expression were normalized by parallel amplification and quantification of GAPDH mRNA levels using forward (5’-ACCACAGTCCATGCCATCAC-3’) and reverse (5’-TCCACCACCCTGTTGCTGTA-3’) primers. Brilliant SYBR Green RT-PCR (Invitrogen) was used in the qPCR mix to amplify the cDNA product. Results were quantified using the ΔΔCT method.

### Live cell imaging

HUVECs (1×10^4^ cells per 200μL media; plate area 34mm^2^) were seeded on Greiner Bio-One CELL view Cell Culture Slides (Fisher Scientific, Pittsburgh, PA) and allowed to adhere overnight. Cells were either left untreated in EGM-2 media, treated with 50pg/mLof IFNɣ, or 50pg/mL of IFNɣ plus 25 μM of non-selective lysosome blocker, hydroxychloroquine, in duplicate for each experiment (n=4). Cells were then stained with LysoTracker red (LTR) probes (ThermoFischer Scientific, Waltham, Ma) and MitoTracker green (MTR) probes (ThermoFischer Scientific, Waltham, Ma) for 30 minutes. Media was replaced with serum free RPMI with glutamine (Mediatech Inc., Manassas, VA). When multiple slides were run, plates were staggered to prevent variation due to time elapsed since staining. Fluorescent microscopy was performed with a Nikon Eclipse Ti with a Plan Apo λ 60x/1.4 Oil Ph3 objective, narrow pass filters, and an Andor Zyla sCMOS 5.5 camera operated by Nikon Elements. For lysosome assessments, the fluorescence intensity as measured by the integrated density was scored using ImageJ 1.51 Java 1.8 running on Windows 7 or 10. For mitotracker images, mitochondrial network morphology per cell was assessed using the Mito-Morphology set of macros outfitted for the FIJI distribution of ImageJ as described [29]. The tools and instructions for their usage can be found at https://github.com/ScienceToolkit/MiNA.

### Autophagy assessments

Autophagophore component proteins LC3-II/I were assessed by immunoblot. PVD membranes were treated with rabbit anti-LC3 primary antibodies (1 μg/mL) (Cell Signaling) diluted in 5% BSA/TBS-T followed by HCRP-conjugated anti-rabbit secondary antibodies (1:20000). Fluorescence units were quantified in the context of APOL1 staining using ImageJ 1.51 Java 1.8 running on Windows 7 or 10.

In parallel, single HUVEC cultures of 40,000 cells per 300μL of cell media representing each genotype were plated on 0.1% gelatin-coated cover slips (BD Biosciences) housed in 24-well plates (1.9 cm^2^ per well). HUVECs were either left untreated or given 50pg/mL of IFNɣ. HUVECs representing each genotype were again given 50pg/mL of IFNɣ plus 25 μM of non-selective lysosome blocker, hydroxychloroquine, therefore recapitulating the hypothesized lysosomal defect in variant-carrying cells. After treatment for 18 hours overnight, HUVECs were washed with PBS and fixed with 3.7% formaldehyde in PBS for 10 minutes. The cover slips were again washed with PBS and cells permeabilized with 0.5% Triton in PBS for 20 minutes. Following an additional wash step, cover slips were treated with PBS gelatin solution as a blocking step for 1 hour. They were then stained with a DAPI DNA dye (Vector Laboratories) and primary antibodies to anti-human SQSTM1/p62 (Abcam) (both raised in rabbit) diluted in PBS gelatin solution at concentrations of 1:300 each. Cells were again washed and stained with anti-rabbit TRIT-C and Alexa-488 (Fisher Scientific) diluted in PBS gelatin at concentrations of 1:300. The cover slips were mounted on glass slides for visualization.

Fluorescent microscopy was performed with a Nikon Eclipse Ti with a 60X N.A. 1.40 Plan APO objective, narrow pass filters, and an Andor Zyla sCMOS 5.5 camera operated by Nikon Elements. Puncta were scored using ImageJ 1.51 Java 1.8 running on Windows 7 or 10. Images of cells were taken to make sure full cells were in the field for measurement. No choices were made based on morphology or intensity. All cells that were fully in each field were traced. A macroinstruction was written to locate discrete bright spots which were identified as puncta. A measurement including a 3-pixel-radius circle centered on each punctum was measured and, for each cell, summed. The integrated density of the total puncta per cell was reported.

### Mitochondrial Respirometry Assay

Forty-thousand HUVECs representing each genotype (0RV, 1RV, and 2RV) were seeded on V7 cell culture plates (Seahorse Bioscience). Cells were either left untreated or treated with IFNɣ (50pg/mL) for 18 hours overnight. One hour prior to measurement, cell culture media was replaced with assay media (3mM glucose, 1 mM sodium pyruvate, and 1.5 mM glutamine without FBS at a pH of 7.4). Port injections of oligomycin (1 μM), FCCP (0.25 μM) and rotenone/antimycin (1 μM each) were filled for bioenergetics profiling. Cellular respiration was measured using a Seahorse Bioscience XF 24-3 analyzer.

### Electron Microscopy

Cultured cells were fixed in 2.5% glutaraldehyde and 2% paraformaldehyde in 0.1M sodium cacodylate buffer (pH 7.2) for 2 hours at 4°C and post-fixed with 1% osmium tetroxide for 1.5 hours at room temperature, then processed in a standard manner and embedded in EMbed 812 (Electron Microscopy Sciences, Hatfield, PA). Ultrathin sections (60 nm) were cut, mounted on copper grids and stained with uranyl acetate and lead citrate by standard methods. Stained grids were examined under a Talos120C electron microscope and photographed with a Gatan OneView camera. Twenty random cells in each sample were imaged for morphological analysis.

### Statistical Analysis

For each genotype and experimental condition, data were expressed as mean ± standard deviation. Medians were used when the population cannot be assumed to be normally distributed. Two sample t-tests for two-group comparisons and ANOVA for multiple-group comparisons were used. Data normality was assessed by visual examination of the observed distributions and Kolmogorov-Smirnov tests. Equality of variance was assessed by F-tests. mRNA expressions and cell densities were log-transformed to better satisfy normality. If F-tests failed to reject the hypothesis of unequal variances, two sample t-tests with equal variances for two-group comparisons and ANOVA for multiple-group comparisons were used instead. When ANOVA tests rejected the null hypothesis, post hoc pairwise comparisons were performed. All statistics were carried out using IBM SPSS software. The level of significance was set at 0.05.

## Results

A unique resource, donor Human Umbilical Vein Endothelial Cells (HUVECS), were isolated from healthy subject umbilical cords representing each of the APOL1 genotypes.

### Inflammatory stimuli increase HUVEC APOL1 Expression

To investigate the intersection between inflammation and APOL1 expression, healthy HUVEC cultures were established, which were isolated and expanded. All APOL1 genotypes were represented as detailed in the methods section. For each experiment, the number of individual HUVEC donors is outlined in S1 Fig. Previous reports suggest that various inflammatory stimuli can increase APOL1 expression in human podocyte and embryonic kidney cell culture models [5]. The capacity of SLE characteristic cytokines including IFNα, IFNɣ, and TNFα to stimulate gene expression in HUVECs was tested. Exposing HUVECs to IFNα, IFNɣ, and TNFα resulted in increased APOL1 expression of 8.7±1.7, 20.8±13.7, and 7.8±2.6 fold respectively, versus untreated HUVECs (for each cytokine vs untreated, p<0.05) (Fig. 1A). Thus multiple inflammatory cytokines that are integral to autoimmune disease were shown to increase APOL1 expression. On immunoblot, IFNɣ-treated HUVECs increased APOL1 protein expression 15.2-fold compared to untreated HUVECs (p=0.01; S1 Fig.). To expose endothelial cells to several circulating cytokines, sera from SLE patients (N=5) and controls were incubated with HUVECs across genotypes. Subsequently, APOL1 expression was assessed. In response to SLE sera, APOL1 expression increased on average 39.8±9.3-fold compared to 3.6±0.7-fold in healthy control sera (Fig 1B; note that for the use of sera derivatives, patient donor demographics and clinical data are shown in Table 1). This increased expression was apparent across genotype (S1 Fig). The genotype of the PCR products was concordant with chromosomal DNA with heterozygous HUVECs expressing both variant and ancestral APOL1 alleles. This result empowered a series of interrogations directed at the consequences of cytokine treatment in heterozygous HUVECs.

**Fig. 1.**
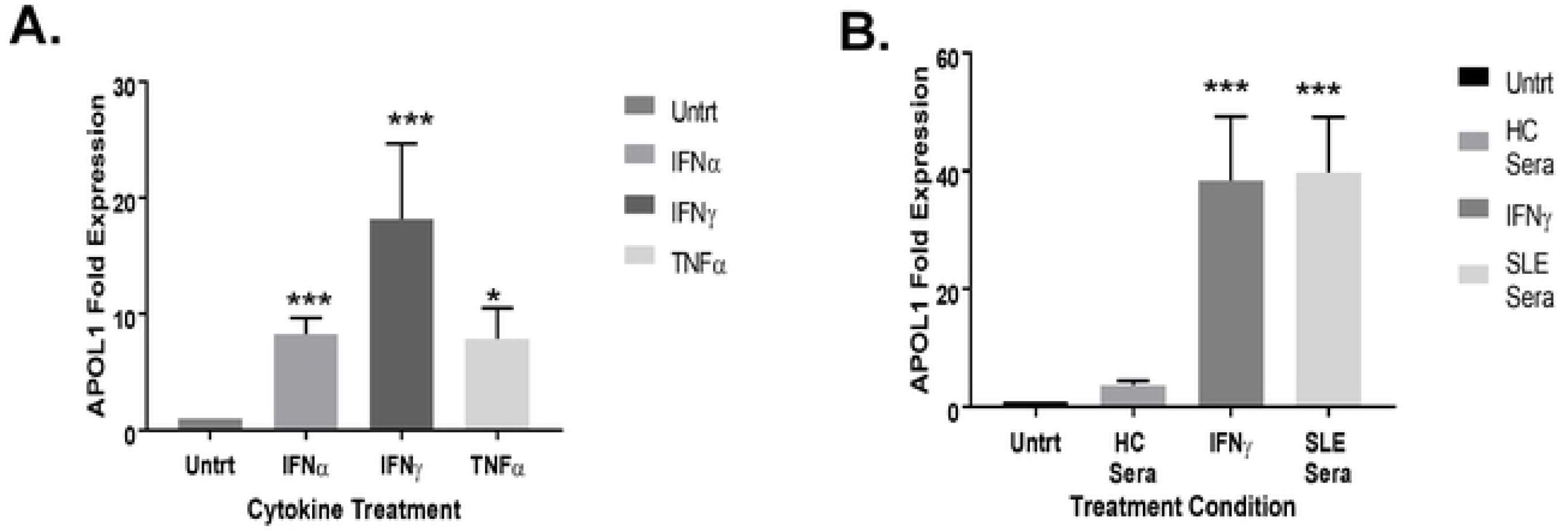
Endothelial cells treated with inflammatory cytokines induce APOL1 expression. A. The upregulation of HUVEC APOL1 transcript in untreated HUVECs compared to IFNα (50pg/mL), IFNɣ (50pg/mL), and TNFα (10ng) treated for 18 hours (average of 5 experiments, 9 HUVEC donors). Shown on the y-axis are 2- ΔΔCT (transcript normalized to GAPDH) values, and shown on the x-axis are cytokine treatment. B. Exposure of HUVECs to sera at 1:1 dilution for 18 hours resulted in an upregulation of APOL1 transcription (average of 5 experiments, 9 HUVEC donors). Shown on the y-axis are 2- ΔΔCT (transcript normalized to GAPDH) values, and shown on the x-axis are treatment conditions. Comparisons are made between the mean fold expression in untreated vs the treatment condition using Kruskal-Wallis test (both 1A and 1B p<0.001) followed by post hoc Dunn test (*** indicates p<0.001, * indicates p<0.05). Abbreviations: Untrt= untreated condition, IFNα= interferon alpha treatment, IFNγ = interferon gamma treatment, TNFα = tumor necrosis factor alpha treatment, HC Sera= healthy control sera, and Systemic Lupus Erythematosus Sera= SLE Sera, each lupus sera (subjects in table 1) was run in triplicate.

**Table 1.**
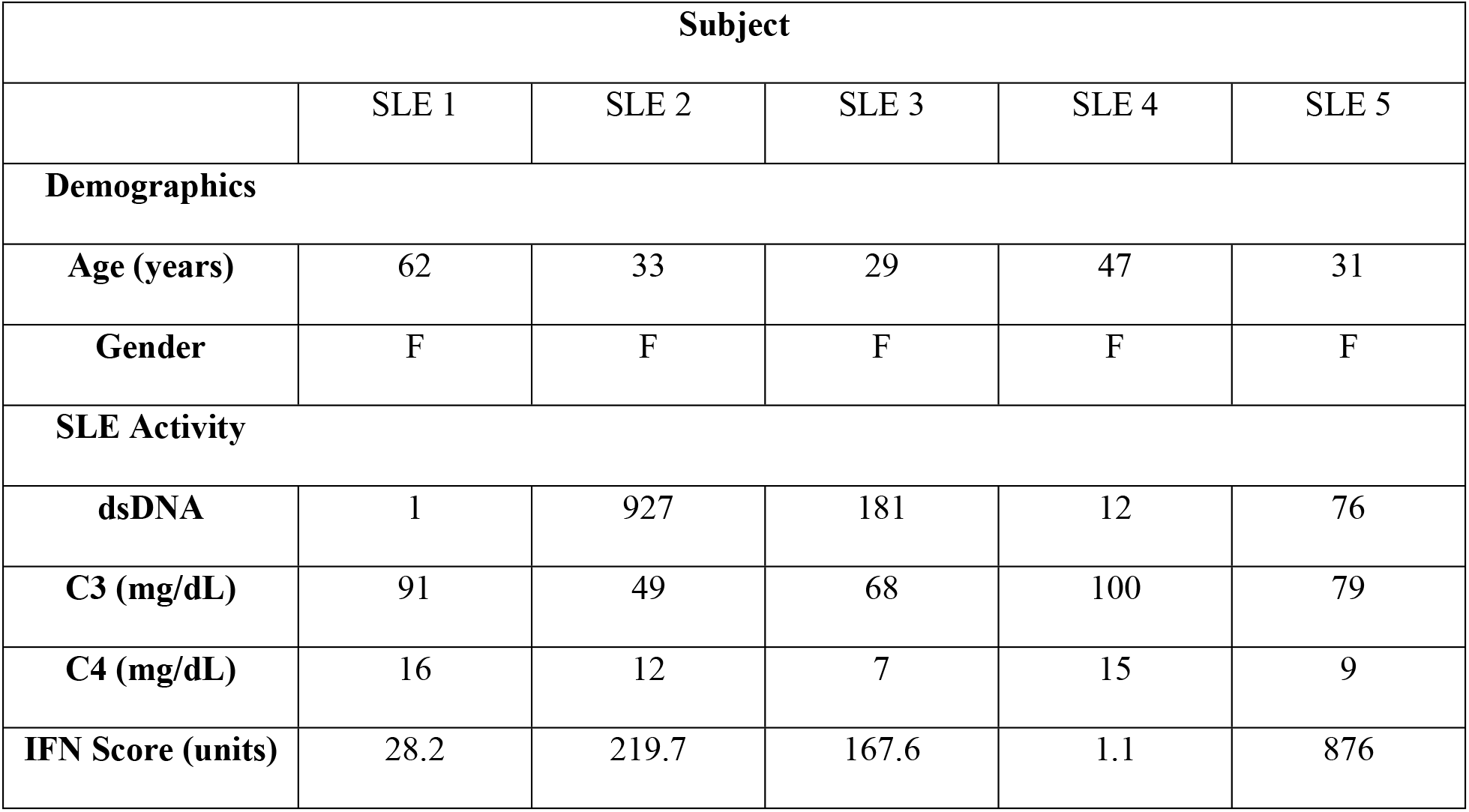
African American (0RV genotype) SLE Sera Donor Demographics, SLE Activity, and IFN score. Demographics and clinical characteristics of SLE serum donor subjects at the time of blood draw. All subjects were African American and APOL1 ancestral allele homozygous.

### APOL1 variant-carrying HUVECs exhibit defects in mitochondrial respiration

While HUVEC IFNɣ exposure results in pleiotropic responses regardless of APOL1 genotype, we were uniquely positioned to examine the consequences of IFNɣ induced expression of APOL1 on a live-cell metabolic assay.

### Live-cell metabolic assays and the stress response of all APOL1 genotypes

Here, the Seahorse XF24 Analyzer was utilized to measure the APOL1 genotype effect on HUVEC mitochondrial respiration. This assay measures cellular mitochondrial function in real time using well-defined inhibitors, oligomycin, FCCP, and Antimycin A [30, 31]. Baseline oxygen consumption rate (OCR) is first measured, followed by the addition of the ATPase inhibitor, oligomycin, in order to evaluate the non-ATPase-dependent OCR. Next an uncoupling agent, FCCP, is added to allow uninhibited electron crossing at the inner mitochondrial membrane, thus measuring the maximum oxygen consumption. Last, a mixture of complex I and complex III inhibitors, rotenone and antimycin A, is added to inhibit mitochondrial respiration completely. The resultant OCR value reflects non-mitochondrial respiration. These measurements can be used to assess overall bioenergetic health index (BHI), which is proportional to reserve capacity and ATP synthase-dependent OCR and inversely proportional to proton leak and non-mitochondrial OCR [30].

We measured OCR in each of the genotypes by treatment condition. As shown in Fig 2 A-C, at baseline, OCR was higher in the 0 risk variant (RV) HUVECs than the 1 or 2 RV HUVECs (89.9±5.6 pmol/min vs 71.7±4.5 pmol/min vs 66.5±3.2 pmol/min respectively; p=0.002). Likewise, maximum OCR was higher in 0 RV HUVECs; values dropped with each RV copy, with means of 152.7±10.7 pmol/min in 0 RV carriers, 122.3±9.6 pmol/min in 1 RV carriers, and 102.6±4.6 pmol/min in 2 RV carriers (p=0.001). With the addition of IFNɣ, maximum OCR fell to 133.2±11.3 pmol/min, 100.8±6.2 pmol/min, and 92.9±6.1 pmol/min in the 0, 1, and 2 RV-carrying HUVECs respectively (p=0.002). This reduction trended toward significance in IFNɣ-treated 1 RV HUVECs (p=0.06).

**Fig 2.**
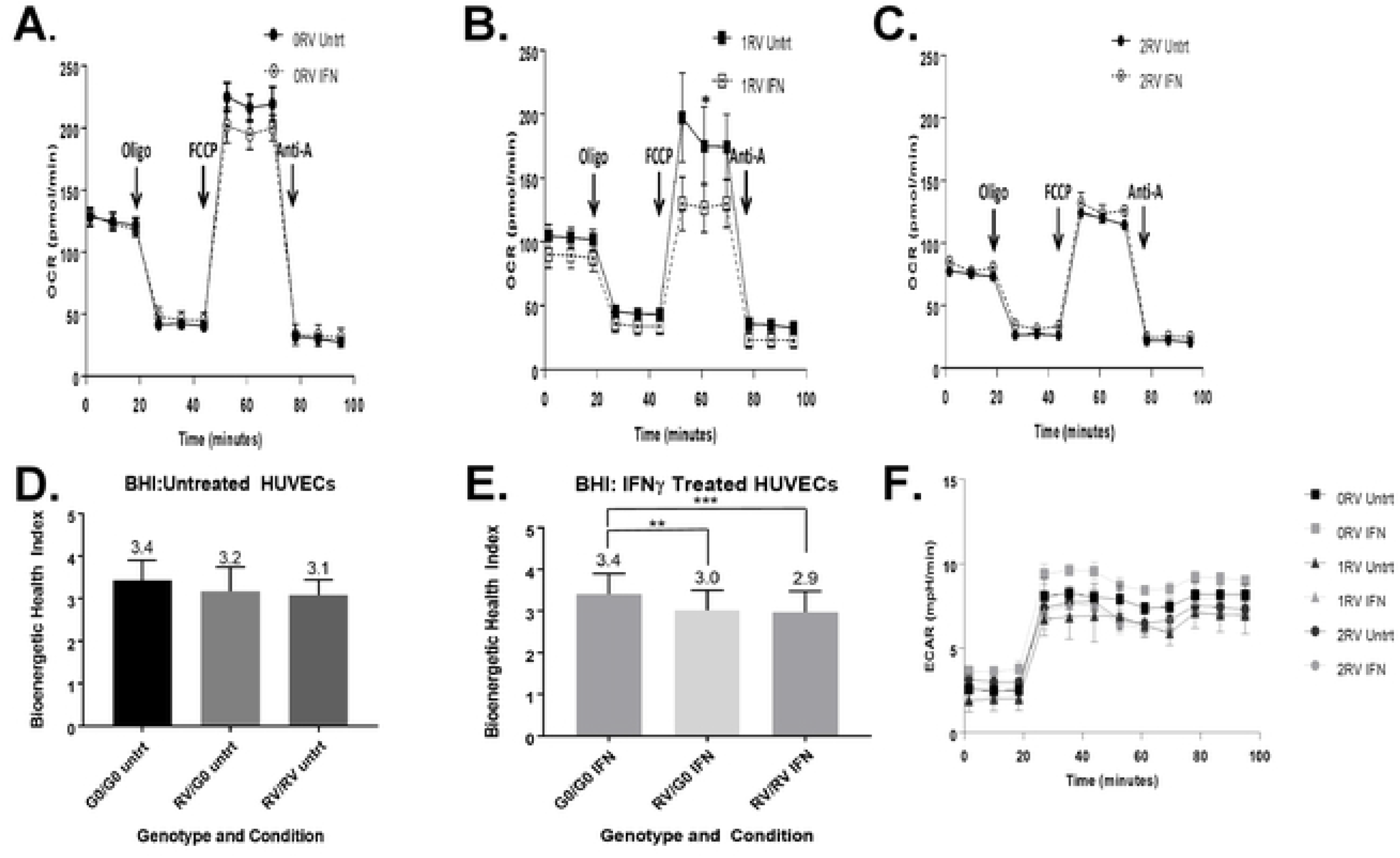
The bioenergetic profiles of HUVECs across genotype. APOL1 risk variant associations with attenuated mitochondrial function including maximum respiration, reserve respiration capacity, glycolytic capacity, and bioenergetic health index. Live HUVEC metabolic assays were performed using the Seahorse XF platform. Genotypes are represented as follows: 0RV, 1RV, and 2RV (RV= risk variant). Treatment conditions included: no treatment (Untrt) or stimulation using IFN (50pg/mL, 18h overnight). A-C. In this assay, oxygen consumption rate (OCR) was measured at baseline, upon oligomycin (oligo), FCCP, and antimycin A treatments. Representative oxygen consumption profiles are shown. D-E. Bioenergetic health index (BHI) calculated by APOL1 genotype and treatment condition (5 experiments averaged representing 9 HUVEC donors). F. Representative raw tracings of extracellular acidification rate (ECAR) after addition of Oligomycin by genotype and condition. P-values were calculated using one-way analysis of means for cross genotype comparisons. Where the three-comparison ANOVA was significant, post-hoc two comparison analysis was completed. * indicates P<0.05, ** indicates p<0.01, and *** indicates p<0.001.

As shown in Fig 2 D-E, there were no statistically significant differences in bioenergetic health index (BHI) in untreated HUVECs across genotype (0 RV=3.43±0.47, 1 RV=3.20±0.57, 2 RV=3.10±0.37; p=0.06); however, treating HUVECs with IFNɣ decreased BHI in 1 and 2 RV-carrying cells resulting in significant differences among genotype groups (0 RV=3.41±0.49, 1 RV=3.02±0.48, 2 RV=2.97 ± 0.51; p=0.025). Results by experiment and sample are shown in S2 Fig. While this deficiency was apparent in both treated and untreated 2 RV-carrying cells, 1 RV-carrying HUVECs exhibited a difference only after IFNɣ treatment, again suggesting that in these cells the adverse phenotype is inducible.

To further determine the RV effect on metabolic capacity in HUVECs, the Seahorse XF glycolysis test was used to measure extracellular acidification rate (ECAR). Glycolysis is an oxygen independent ATP production process that converts glucose to lactate. Lactate is the major source of free protons in the culture medium [32]. Therefore, by measuring the rate at which the extracellular medium becomes acidified, the XF24 analyzer measures the glycolytic rate in real time. With the addition of stressor, oligomycin, which blocks mitochondrial respiration, cells are forced to utilize oxygen-independent glycolysis to meet energy demands [32]. The glycolytic capacity can therefore be measured by the difference between baseline and oligomycin-treated (stressed) glycolytic rate [32]. Overall the glyocolytic capacity in untreated endothelial cells was significantly lower in 2 RV carriers than in 1 or 0 RV carriers (0 RV: 5.4 mph/min, 1 RV: 4.8 mph/min, 2 RV: 4.2 mph/min, p=0.02). This parameter was lowered by IFNγ treatment (0RV: 4.7 mph/min, 1RV: 4.5 mph/min, 2RV: 3.9 mph/min, p = 0.04, Fig 2F).

Consistent with differences in OCR and ECAR, ATP production varied across genotype and treatment condition. At baseline, ATP production was numerically, but not significantly higher – 45.6pmol/min – in 0 RV carriers, compared to 33.2pmol/min and 34.2pmol/min in 1 or 2 RV carriers (p=0.09). IFNγ treatment increased ATP production in each genotype, however to a lesser extent in variant carriers. The resepective values in the 0, 1, and 2 RV HUVECs were 48.7pmol/min, 34.7pmol/min, and 37.7pmol/min (p=0.02). These results support an association between APOL1 risk variants and impaired mitochondrial function characterized by reduced maximum respiration, reserve respiration capacity, glycolytic capacity, and bioenergetic health index. HUVECs carrying the variants exhibited a more senescent phenotype with overall lower energy production. This observation was more pronounced with each RV copy (i.e. 2 RV > 1 RV > 0 RV), and was exacerbated by IFNγ treatment particularly in the 1RV HUVECs.

## Mitochondrial structure

To determine genotype-associated differences in mitochondrial structure, mitochondria were stained with fluorescent dye, MitoTracker. HUVECs of each genotype were either left untreated, or given IFNɣ overnight and evaluations of mitochondrial ultrastructure included indirect (use of fluorescent dye, MitoTracker), and direct (transmission electron microscopy (TEM)). For the former, sets of 10 HUVEC images across genotype donors and treatment conditions were evaluated for mitochondrial morphology per cell including length and number of branches using an automated protocol [29] (representative images shown in Fig 3 A-B). In untreated HUVECs, 2RV carriers exhibited shorter branch length (sum branch length: 0RV: 4.3μm 1RV: 3.3μm 2RV: 3.2μm; p= 0.17) and less networking (mean networked branches: 0RV: 2.0 1RV: 2.1 2RV: 1.8; p= 0.6) though these differences did not reach statistical significance (Fig 3 C-D). Upon treatment with IFNγ, genotype-associated differences became significant (sum branch length: 0RV: 4.6μm 1RV: 3.1μm 2RV: 2.7μm; three group comparison p=0.003; Post hoc 2 group comparison 0RV vs 1RV p=0.03; 0RV vs 2RV p=0.004. Mean networked branches: 0RV: 2.3 1RV: 1.8 2RV: 1.7; three group comparison p= 0.001. Post hoc 2 group comparison 0RV vs 1RV p=0.01; 0RV vs 2RV p=0.002; Fig 3 C-D). TEM images were concordant with this finding. Untreated cells across genotype showed small differences in mitochondrial area (median area for 0, 1, and 2 allele HUVECs=0.09μm^2^, 0.08μm^2^, 0.07μm^2^ respectively) (S3 Fig). Upon IFNγ treatment, these differences became more pronounced (median area + IFNɣ 0 RV=0.11μm; 1 RV=0.10μm; 2 RV=0.07 p <0.001) (S3 Fig).

**Fig 3.**
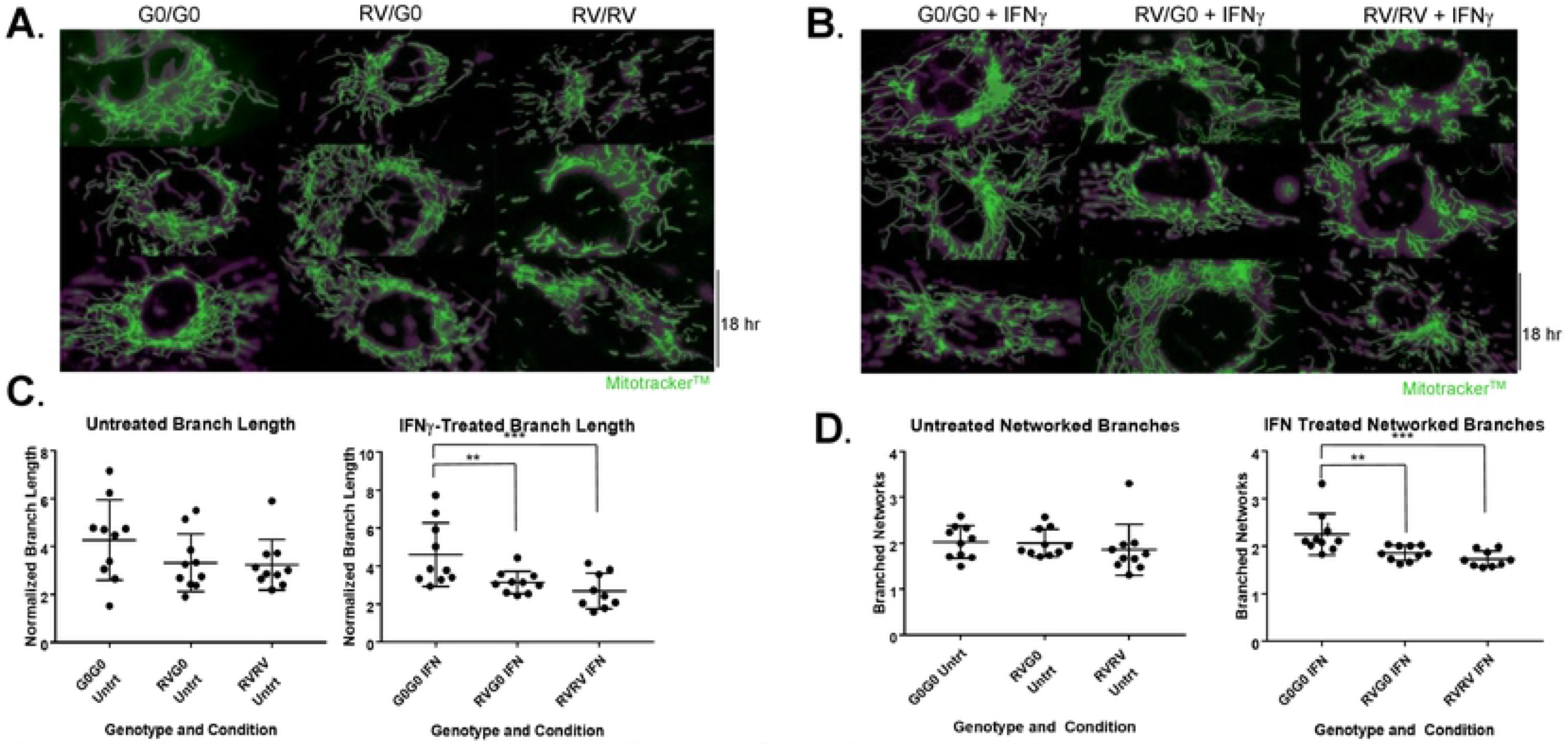
Assessment of mitochondrial structure in resting and stimulated HUVECs using MitoTracker, a fluorescent proxy of ultrastructure. Mitochondrial structure including branch length and networking is attenuated in HUVECs with 1RV or 2RV versus 0RV APOL1 genotype both at baseline and with IFNγ 50pg/mL-treatment. Human umbilical vein endothelial cells representing each genotype, 0 risk variant (G0/G0 left columns), 1 risk variant (RV/G0 middle columns), and 2 risk variant (RV/RV right columns) were either left untreated (A) or treated with IFNɣ 50pg/mL (B) for 18 hours overnight. Representative immunofluorescence (MitoTracker green stained) images are shown (overall experiment, 10 cells per genotype and condition were measured. In total 12 HUVEC donors and 4 experiments were averaged). C-D Mitochondrial branch length (C) and networked branches (D) measured on confocal microscopy versus genotype and treatment condition (x axis) using the Mitochondrial Network Analysis (MiNA) tools available in the FIJI distribution of ImageJ. Mitochondrial length was measured in microns (μM). Each additional risk variant associated with a reduced degree of mitochondrial networking--an effect that became statistically significant across the genotypes upon treatment with IFNɣ. P-values were calculated using Kruskal-Wallis test for cross genotype comparisons, and Wilcoxon rank sum test for comparing untreated group and IFN group. P<0.01 is indicated by **, and p<0.001 is indicated by ***.

### APOL1 variant-Carrying HUVECs exhibit lysosomal defects

Lysosomes serve as a cellular “digestive system,” and their function is highly dependent on both an acidic pH and an ATP-dependent pump. To determine genotype-associated differences in lysosome structure, we stained HUVECs representing each APOL1 genotype and treatment condition with fluorescent dye, Lysotracker. At baseline, 2 RV HUVECs exhibited significantly lower lysotracker staining intensity than 0 or 1 RV HUVECs (p<0.001) (Fig 4A). IFNɣ exposure significantly decreased lysotracker staining intensity in the 0 and 1 RV carriers (p=0.04 and p<0.001 respectively); in the 2RV carriers lysosome staining intensity did not change with IFNγ treatment but remained lower than that of the other two genotypes (Fig 4B). Next, HUVECs were treated with both IFNɣ and HCQ. HCQ is a reagent that blocks lysosome acidification thereby arresting both organelle function and turnover [33, 34]. Treatment of cells with 25 μM of hydroxychloroquine (HCQ) did not influence HUVEC APOL1 expression (S4 Fig). In all conditions, the co-treatment of IFNɣ and HCQ resulted in increased lysosome staining intensity (Fig 4C). This effect, however, was significantly less in both 1 and 2 RV-carrying cells. Lysosome staining intensity is quantified in Fig 4D, and results by experiment and sample are shown in S5 Fig. Taken together, each additional RV copy associated with less HUVEC lysosome staining intensity—an observation that was exacerbated by IFNγ treatment.

**Fig 4.**
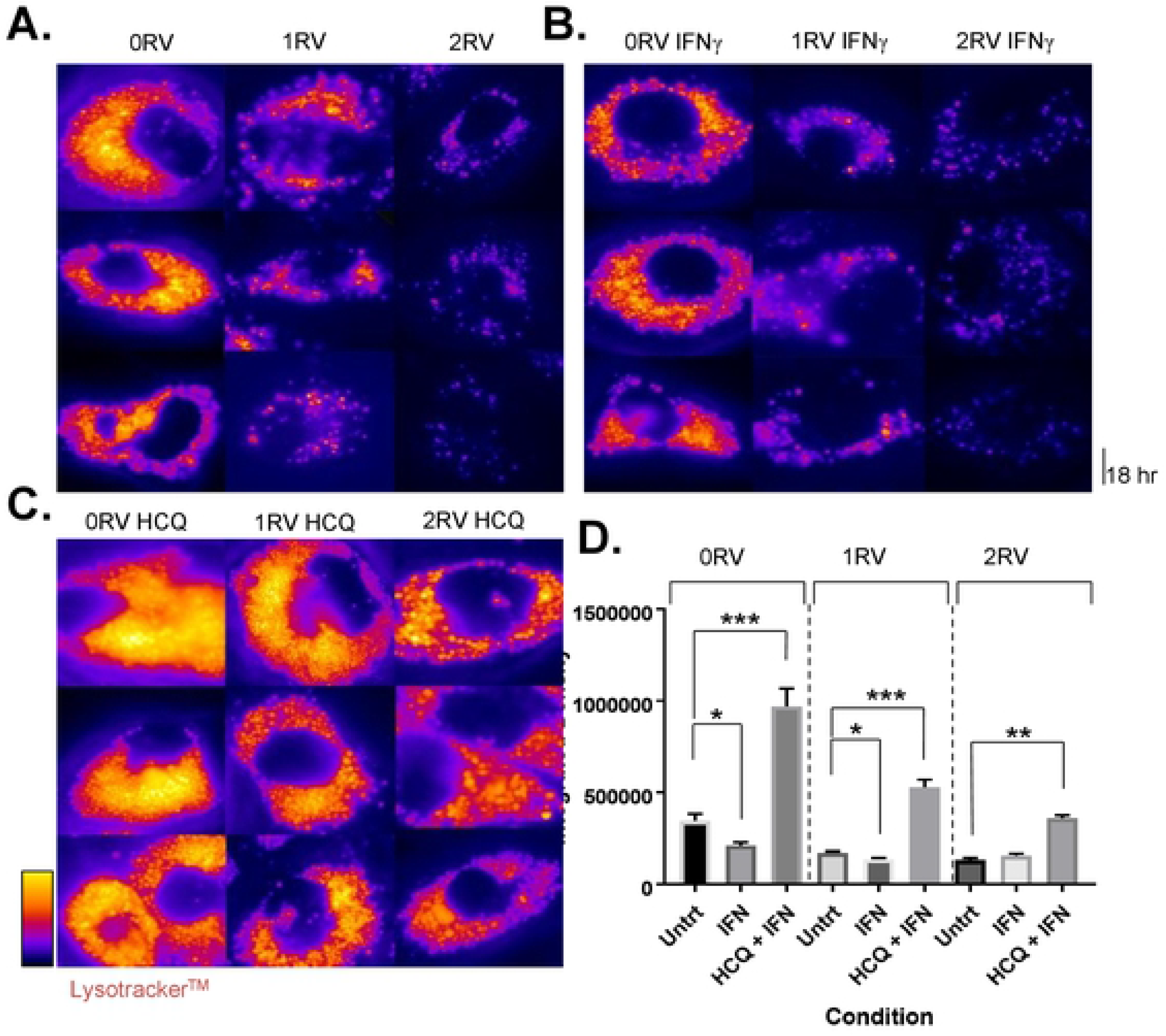
Assessment of lysosomal structure in resting and stimulated HUVECs with or without hydroxychloroquine (HCQ) using LysoTracker. The presence of 1RV or 2RVs associated with less lysosome staining by lysotracker at baseline or with IFNγ treatment. Preventing lysosome turnover with the addition of HCQ treatment increased lysosome staining to a lesser degree in RV carrying HUVECs. A. Immunofluorescent images of untreated HUVECs representing each genotype, 0RV (left column), 1RV, and 2RV (right column). B. Representative images of IFNɣ 50pg/mL-treated HUVECs across genotype (parallel layout as per A) for 18 hours overnight. C. Representative images of HCQ (25μM) plus IFNɣ-treated HUVECs across genotype (layout as per A) LysoTracker staining was performed. Images were captured by florescent microscopy using a Nikon Eclipse Ti and representative microphotographs were selected. D. The average lysosome intensity per region of interest (Integrated Density) for each genotype and treatment condition group. P-values were calculated using Kruskal-Wallis test for cross genotype comparisons, and Wilcoxon rank sum test for comparing untreated group to the IFNγ treated or HCQ plus IFNγ group.*P<0.05; **p<0.01, ***p<0.001.

### APOL1 variant-carrying HUVECs display defects in autophagic flux

Autophagosome maturation and degradation (flux) is contingent upon a functioning lysosome [35] which was demonstrated to be compromised in APOL1 variant-carrying HUVECs (above). Therefore, autophagosomes were evaluated using both fluorescent microscopy of SQSTM1 (p62) and transmission electron microscopy (TEM). SQSTM1 (p62) is an autophagophore shuttle protein that is degraded through autophagy and has been utilized in the literature to measure autophagic flux on microscopy. HUVECs were stained for p62 and the number of autophagic puncta (log transformed) per cell was observed. HCQ was again used to arrest the degradation of the autophagosomes by blocking lysosome acidification [36]. Therefore, a comparison of APOL1 genotype-dependent differences in autophagy at baseline, upon IFNɣ treatment, and IFNɣ plus HCQ treatment (autophagic flux inhibition) was established. To assess the degree to which autophagic flux was impaired, the puncta count at baseline or upon IFNɣ treatment was compared to the HCQ-treated condition.

It was observed that in untreated HUVECs, autophagosome count was the lowest in 0 RV cells and increased with each additional variant allele (log autophagosome count per genotype: 0RV: 1.1±0.57; 1RV: 1.6±0.48; 2RV: 2.0±0.70, p<0.001 Fig 5A). Across genotypes, IFNɣ exposure increased autophagosome count (0 RV: 1.3±0.45, 1 RV: 1.8±0.44, 2 RV: 2.2±0.31, p<0.001 Fig 5B). In each genotype group, the number of autophagosomes increased in the IFNɣ plus HCQ–treated condition; however 2 RV HUVECs exhibited the highest autophagosome count (0 RV: 1.7±0.38, 1 RV: 2.0±0.5, 2 RV: 2.5±0.55, p<0.001 Fig 5C). Autophagosomes were confirmed on TEM by genotype and treatment condition (Fig 5D). Microscopy results are quantified in Fig 5E, and confirmed by p62 immunoblot in Fig 5F. Results by experiment are shown in S6 Fig. A supporting immunoblot of the LC3 II/I ratio by genotype and treatment condition is shown in S7 Fig. Taken together, these results support an association between HUVEC APOL1 genotype and autophagic flux inhibition. Congruent with the lysosome staining results, each additional risk variant had an effect on autophagic flux with interferon exacerbating the phenotype–particularly in the heterozygous condition.

**Fig 5.**
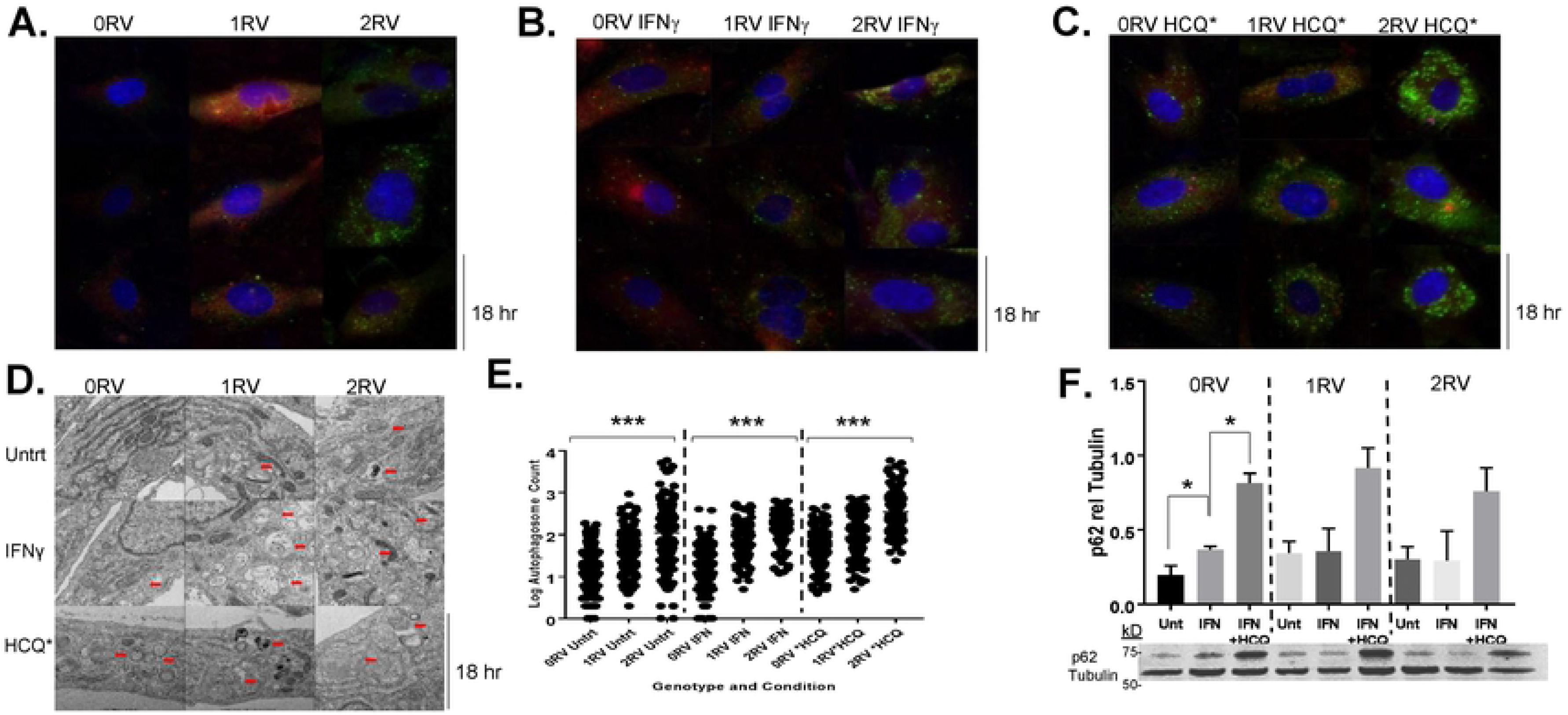
APOL1 risk variant-carrying HUVECs display autophagic flux deficiencies. Assessment of autophagosome accumulation using SQSTM1 (p62) staining, a proxy for autophagic flux inhibition. A. Representative immunofluorescence images of untreated HUVECs stained for SQSTM1 (p62) across APOL1 genotype: 0 risk variants (left column), 1 risk variant (middle column), 2 risk variants (right column). B. Representative immunofluorescence images of IFNɣ (50pg/mL)-treated HUVECs stained for SQSTM1 (p62) across APOL1 genotype C. Representative immunofluorescence images of IFNɣ plus HCQ (25μM)-treated HUVECs stained for SQSTM1 (p62) across APOL1 genotype D. Autophagosomes were confirmed by transmission electron microscopy (Columns: 0RV left, 1RV middle, 2RV right; Rows: Untrt=untreated top, IFNɣ-treated middle, IFNɣ plus HCQ-treated bottom). E. Log transformed values of p62 positive puncta per cell (y axis) and the treatment condition (x axis) are shown. HUVEC genotype is labeled from left to right. F. Immunoblot of HUVEC lysates showing SQSTM1 (p62) protein concentration compared to tubulin loading control by genotype and treatment condition. Note: HUVECs were treated for 18 hours. *P<0.05; ***P<0.001 were calculated from one-way analysis of means for cross genotype comparisons. Where ANOVA rejected the null hypothesis, a post-hoc 2 group analysis was performed.

## Discussion

Overall, these data support functionally relevant associations between APOL1 risk genotypes and various endothelial organelle readouts. HUVECs exposed to cytokines characteristically detected in SLE patients, IFNα, IFNɣ, and TNFα, strongly up-regulated APOL1 gene expression. Exploiting the use of HUVECs expressing ancestral and variant APOL1 alleles, it was demonstrated for the first time that variant expression associates with functional consequences such as decreased mitochondrial metabolic potential and mitochondrial fragmentation. Specifically, in 2RV and IFNɣ-treated RV heterozygous HUVECs, maximum respiration, reserve respiration capacity, glycolytic capacity, ATP production, and bioenergetic health index were attenuated relative to the ancestral allele homozygous HUVECs. Using a parallel set of conditions, the presence of 1RV or 2RVs also associated with decreased lysosome staining by lysotracker as well as an inhibition of autophagocytic flux, which was based on fluorescent evaluations of SQSTM1 and confirmed by transmission electron microscopy. Taken together, these data support that 2 RVs—and under inflammatory stress 1 RV--contribute to a senescent endothelial cell phenotype, which is characterized by overall lower energy production and untoward consequences to autophagosome maturation and degradation (flux), an event that is contingent upon a functioning lysosome. Given the prior reporting of vascular disease in APOL1 variant-carrying SLE patients, these observations in HUVECs may offer mechanistic insights [4, 21, 37].

These data add to an overall understanding of APOL1 variant effects in endothelial cells, and the interacting microenvironment which may potentiate cellular injury. Although traditionally this has been characterized by defective nitric oxide synthesis, recent work has demonstrated that autophagy deficiencies contribute to atherogenesis [37, 38]. These results demonstrate a potential mechanistic link between SLE-associated inflammation and APOL1 risk variant status via inducible endothelial injury.

APOL1 is a five-domain protein with several intracellular functions [6, 16, 18, 39]. Its expression is up-regulated by cellular stress including inflammatory signals, nutrient deprivation, and hypoxia [9, 10]. APOL1’s colicin-like pore forming domain may be inserted into cell membrane, lysosome, or mitochondrial phospholipid bilayers in a pH dependent fashion [40]. The G1 and G2 mutations allow for pore formation at lower levels of APOL1 gene expression [40]. Mitochondria may be injured by APOL1 variants either directly, or indirectly due to defects in lysosomes and autophagic flux. APOL1 has been shown to cause toxicity by disrupting lysosomes in human podocytes [15]. Since intracellular organelles are not viewed as static and in isolation, there could be a potential consequence to mitochondria. For example, it has been shown that lysosome membrane permeability allows escape of lysosome hydrolases, including cathepsin-B, which mediate mitochondrial outer membrane permeability and loss of inner-membrane potential [41]. Future studies exploring these mechanisms in APOL1 risk variant-carrying tissues could better explain gene-associated pathobiology. Also mitophagy, an autophagy analogue, is the predominant means by which cells dispose of energy-depleted mitochondria [42]. Finally, others have shown that the APOL1 pore co-localizes with the mitochondria in human embryonic kidney cells, directly causing membrane permeability [43]. The notion that these processes are active in APOL1-expressing endothelial cells is a novel and potentially biologically relevant finding.

While there are current APOL1 primary cell culture models in the literature, most have employed viral vector systems to deliver the gene [5, 12, 44]. Importantly, these models showed that APOL1 variant expression coincides with mitochondrial fragmentation, lysosome compromise, and an abundance of autophagosomes [13, 15]. However, exaggerated APOL1 expression beyond that expected in native cells poses a limitation on the clinical interpretation of these results. Moreover, viruses themselves can not only induce APOL1 expression but also engage other autophagy pathways, making it difficult to assign risk variant-mediated effects [45] Therefore, HUVECs naturally expressing the variants were utilized acknowledging shortcomings inherent in this approach as well.

Though experiments were repeated across multiple donors of each genotype, the possibility that non-APOL1-related genetic variation contributed to our findings cannot be excluded. Both male and female samples were utilized, and despite even distribution across genotype, differences in autophagy due to sex cannot be excluded. Also, HUVEC cells may not recapitulate endothelial cell behavior in other vascular beds more relevant to atherosclerotic disease. Finally, the threshold at which APOL1 expression influences lysosome function is not clear based on this study. Further work determining the intracellular concentration of APOL1 and timing of lysosome injury is warranted given the above findings.

In sum, variant APOL1 expression, particularly in the presence of inflammatory stimuli typical of autoimmune diseases such as SLE, confers endothelial dysfunction mirrored in mitochondrial stress, lysosomal dysfunction and impaired autophagic flux, Treatments aimed at compensating for these metabolically compromised cellular states may improve the vascular consequences facing APOL1 variant carriers.

## Acknowledgements

The authors would like to thank the NYU Langone Department of Obstetrics and Gynecology for aiding in the curation of HUVEC samples, and the NYU CTSI for providing support in conceptualizing this project. All imaging was completed in the NYU DART Microscopy Core.

## Supporting Information

**S1 Fig. This is the S1 Fig title**. This is the S1 Fig legend.

**S2 Fig. This is the S2 Fig title**. This is the S2 Fig legend.

## References

1. Genovese G, Friedman DJ, Ross MD, Lecordier L, Uzureau P, Freedman BI, et al. Association of trypanolytic ApoL1 variants with kidney disease in African Americans. Science. 2010;329(5993):841–5. doi: 10.1126/science.1193032. PubMed PMID: 20647424; PubMed Central PMCID: PMC2980843.

2. Ko WY, Rajan P, Gomez F, Scheinfeldt L, An P, Winkler CA, et al. Identifying Darwinian selection acting on different human APOL1 variants among diverse African populations. Am J Hum Genet. 2013;93(1):54–66. Epub 2013/06/19. doi: 10.1016/j.ajhg.2013.05.014. PubMed PMID: 23768513; PubMed Central PMCID: PMCPMC3710747.

3. Friedman DJ, Kozlitina J, Genovese G, Jog P, Pollak MR. Population-based risk assessment of APOL1 on renal disease. J Am Soc Nephrol. 2011;22(11):2098–105. doi: 10.1681/ASN.2011050519. PubMed PMID: 21997396; PubMed Central PMCID: PMC3231785.

4. Blazer A, Wang B, Simpson D, Kirchhoff T, Heffron S, Clancy RM, et al. Apolipoprotein L1 risk variants associate with prevalent atherosclerotic disease in African American systemic lupus erythematosus patients. PLoS One. 2017;12(8):e0182483. Epub 2017/08/30. doi: 10.1371/journal.pone.0182483. PubMed PMID: 28850570; PubMed Central PMCID: PMCPMC5574561.

5. Nichols B, Jog P, Lee JH, Blackler D, Wilmot M, D’Agati V, et al. Innate immunity pathways regulate the nephropathy gene Apolipoprotein L1. Kidney Int. 2015;87(2):332–42. doi: 10.1038/ki.2014.270. PubMed PMID: 25100047; PubMed Central PMCID: PMC4312530.

6. O’Toole JF, Bruggeman LA, Madhavan S, Sedor JR. The Cell Biology of APOL1. Seminars in nephrology. 2017;37(6):538–45. Epub 2017/11/08. doi: 10.1016/j.semnephrol.2017.07.007. PubMed PMID: 29110761; PubMed Central PMCID: PMCPMC5678957.

7. Kent WJ, Sugnet CW, Furey TS, Roskin KM, Pringle TH, Zahler AM, et al. The human genome browser at UCSC. Genome Res. 2002;12(6):996–1006. Epub 2002/06/05. doi: 10.1101/gr.229102. PubMed PMID: 12045153; PubMed Central PMCID: PMCPMC186604.

8. Langefeld CD, Comeau ME, Ng MCY, Guan M, Dimitrov L, Mudgal P, et al. Genome-wide association studies suggest that APOL1-environment interactions more likely trigger kidney disease in African Americans with nondiabetic nephropathy than strong APOL1-second gene interactions. Kidney Int. 2018. Epub 2018/06/11. doi: 10.1016/j.kint.2018.03.017. PubMed PMID: 29885931.

9. Vanhollebeke B, Pays E. The function of apolipoproteins L. Cellular and molecular life sciences : CMLS. 2006;63(17):1937–44. doi: 10.1007/s00018-006-6091-x. PubMed PMID: 16847577.

10. Wan G, Zhaorigetu S, Liu Z, Kaini R, Jiang Z, Hu CA. Apolipoprotein L1, a novel Bcl-2 homology domain 3-only lipid-binding protein, induces autophagic cell death. J Biol Chem. 2008;283(31):21540–9. doi: 10.1074/jbc.M800214200. PubMed PMID: 18505729; PubMed Central PMCID: PMC2490785.

11. Blazer A, Chang M, Robins K, Buyon J, Clancy R. Macrophages, APOL1 Genotype, & Immunometabolism in CVD (MAGIC). Journal of Clinical and Translational Science. 2019;3(s1):1. Epub 26 March 2019. doi: 10.1017/cts.2019.339.

12. Olabisi OA, Zhang JY, VerPlank L, Zahler N, DiBartolo S, 3rd, Heneghan JF, et al. APOL1 kidney disease risk variants cause cytotoxicity by depleting cellular potassium and inducing stress-activated protein kinases. Proc Natl Acad Sci U S A. 2016;113(4):830–7. doi: 10.1073/pnas.1522913113. PubMed PMID: 26699492; PubMed Central PMCID: PMC4743809.

13. Granado D, Muller D, Krausel V, Kruzel-Davila E, Schuberth C, Eschborn M, et al. Intracellular APOL1 Risk Variants Cause Cytotoxicity Accompanied by Energy Depletion. J Am Soc Nephrol. 2017;28(11):3227–38. Epub 2017/07/12. doi: 10.1681/ASN.2016111220. PubMed PMID: 28696248; PubMed Central PMCID: PMCPMC5661279.

14. Beckerman P, Bi-Karchin J, Park AS, Qiu C, Dummer PD, Soomro I, et al. Transgenic expression of human APOL1 risk variants in podocytes induces kidney disease in mice. Nature medicine. 2017;23(4):429–38. doi: 10.1038/nm.4287. PubMed PMID: 28218918.

15. Lan X, Jhaveri A, Cheng K, Wen H, Saleem MA, Mathieson PW, et al. APOL1 risk variants enhance podocyte necrosis through compromising lysosomal membrane permeability. Am J Physiol Renal Physiol. 2014;307(3):F326–36. Epub 2014/06/06. doi: 10.1152/ajprenal.00647.2013. PubMed PMID: 24899058; PubMed Central PMCID: PMCPMC4121568.

16. Heneghan JF, Vandorpe DH, Shmukler BE, Giovinazzo JA, Raper J, Friedman DJ, et al. BH3 domain-independent apolipoprotein L1 toxicity rescued by BCL2 prosurvival proteins. Am J Physiol Cell Physiol. 2015;309(5):C332–47. doi: 10.1152/ajpcell.00142.2015. PubMed PMID: 26108665; PubMed Central PMCID: PMCPMC4556898.

17. Bolanos-Garcia VM, Miguel RN. On the structure and function of apolipoproteins: more than a family of lipid-binding proteins. Progress in biophysics and molecular biology. 2003;83(1):47–68. PubMed PMID: 12757750.

18. Sharma AK, Friedman DJ, Pollak MR, Alper SL. Structural characterization of the C-terminal coiled-coil domains of wild-type and kidney disease-associated mutants of apolipoprotein L1. FEBS J. 2016;283(10):1846–62. doi: 10.1111/febs.13706. PubMed PMID: 26945671; PubMed Central PMCID: PMCPMC4879057.

19. Bruggeman LA, O’Toole JF, Sedor JR. APOL1 polymorphisms and kidney disease: loss-of-function or gain-of-function? Am J Physiol Renal Physiol. 2019;316(1):F1–F8. Epub 2018/10/18. doi: 10.1152/ajprenal.00426.2018. PubMed PMID: 30332315; PubMed Central PMCID: PMCPMC6383195.

20. Limou S, Nelson GW, Kopp JB, Winkler CA. APOL1 kidney risk alleles: population genetics and disease associations. Advances in chronic kidney disease. 2014;21(5):426–33. doi: 10.1053/j.ackd.2014.06.005. PubMed PMID: 25168832; PubMed Central PMCID: PMC4157456.

21. Davignon J, Ganz P. Role of endothelial dysfunction in atherosclerosis. Circulation. 2004;109(23 Suppl 1):III27–32. Epub 2004/06/17. doi: 10.1161/01.CIR.0000131515.03336.f8. PubMed PMID: 15198963.

22. Hochberg MC. Updating the American College of Rheumatology revised criteria for the classification of systemic lupus erythematosus. Arthritis Rheum. 1997;40(9):1725. doi: 10.1002/1529-0131(199709)40:9<1725::AID-ART29>3.0.CO;2-Y. PubMed PMID: 9324032.

23. Crampton SP, Davis J, Hughes CC. Isolation of human umbilical vein endothelial cells (HUVEC). J Vis Exp. 2007;(3):183. Epub 2008/11/04. doi: 10.3791/183. PubMed PMID: 18978951; PubMed Central PMCID: PMCPMC2576276.

24. Niewold TB, Adler JE, Glenn SB, Lehman TJ, Harley JB, Crow MK. Age- and sex-related patterns of serum interferon-alpha activity in lupus families. Arthritis and rheumatism. 2008;58(7):2113–9. Epub 2008/06/26. doi: 10.1002/art.23619. PubMed PMID: 18576315; PubMed Central PMCID: PMC2729701.

25. Hua J, Kirou K, Lee C, Crow MK. Functional assay of type I interferon in systemic lupus erythematosus plasma and association with anti-RNA binding protein autoantibodies. Arthritis and rheumatism. 2006;54(6):1906–16. Epub 2006/06/01. doi: 10.1002/art.21890. PubMed PMID: 16736505.

26. Niewold TB, Wu SC, Smith M, Morgan GA, Pachman LM. Familial aggregation of autoimmune disease in juvenile dermatomyositis. Pediatrics. 2011;127(5):e1239–46. Epub 2011/04/20. doi: peds.2010-3022 [pii] 10.1542/peds.2010-3022. PubMed PMID: 21502224; PubMed Central PMCID: PMC3081190.

27. Feng X, Han D, Kilaru BK, Franek BS, Niewold TB, Reder AT. Inhibition of interferon-beta responses in multiple sclerosis immune cells associated with high-dose statins. Arch Neurol. 2012;69(10):1303–9. Epub 2012/07/18. doi: 10.1001/archneurol.2012.465. PubMed PMID: 22801747; PubMed Central PMCID: PMC3910505.

28. Harley ITW, Niewold TB, Stormont RM, Kaufman KM, Glenn SB, Franek BS, et al. The role of genetic variation near interferon-kappa in systemic lupus erythematosus J Biomed Biotechnol. 2010;2010:Article ID 706825.

29. Valente AJ, Maddalena LA, Robb EL, Moradi F, Stuart JA. A simple ImageJ macro tool for analyzing mitochondrial network morphology in mammalian cell culture. Acta Histochem. 2017;119(3):315–26. Epub 2017/03/21. doi: 10.1016/j.acthis.2017.03.001. PubMed PMID: 28314612.

30. Chacko BK, Kramer PA, Ravi S, Benavides GA, Mitchell T, Dranka BP, et al. The Bioenergetic Health Index: a new concept in mitochondrial translational research. Clin Sci (Lond). 2014;127(6):367–73. Epub 2014/06/05. doi: 10.1042/CS20140101. PubMed PMID: 24895057; PubMed Central PMCID: PMCPMC4202728.

31. Son JM, Sarsour EH, Kakkerla Balaraju A, Fussell J, Kalen AL, Wagner BA, et al. Mitofusin 1 and optic atrophy 1 shift metabolism to mitochondrial respiration during aging. Aging Cell. 2017;16(5):1136–45. Epub 2017/08/02. doi: 10.1111/acel.12649. PubMed PMID: 28758339; PubMed Central PMCID: PMCPMC5595680.

32. Mookerjee SA, Nicholls DG, Brand MD. Determining Maximum Glycolytic Capacity Using Extracellular Flux Measurements. PLoS One. 2016;11(3):e0152016. Epub 2016/04/01. doi: 10.1371/journal.pone.0152016. PubMed PMID: 27031845; PubMed Central PMCID: PMCPMC4816457.

33. Steinman RM, Mellman IS, Muller WA, Cohn ZA. Endocytosis and the recycling of plasma membrane. J Cell Biol. 1983;96(1):1–27. Epub 1983/01/01. doi: 10.1083/jcb.96.1.1. PubMed PMID: 6298247; PubMed Central PMCID: PMCPMC2112240.

34. Klionsky DJ, Abdelmohsen K, Abe A, Abedin MJ, Abeliovich H, Acevedo Arozena A, et al. Guidelines for the use and interpretation of assays for monitoring autophagy (3rd edition). Autophagy. 2016;12(1):1–222. Epub 2016/01/23. doi: 10.1080/15548627.2015.1100356. PubMed PMID: 26799652; PubMed Central PMCID: PMCPMC4835977.

35. Liu X, Qin H, Xu J. The role of autophagy in the pathogenesis of systemic lupus erythematosus. Int Immunopharmacol. 2016;40:351–61. doi: 10.1016/j.intimp.2016.09.017. PubMed PMID: 27673477.

36. Mauthe M, Orhon I, Rocchi C, Zhou X, Luhr M, Hijlkema KJ, et al. Chloroquine inhibits autophagic flux by decreasing autophagosome-lysosome fusion. Autophagy. 2018;14(8):1435–55. Epub 2018/06/27. doi: 10.1080/15548627.2018.1474314. PubMed PMID: 29940786; PubMed Central PMCID: PMCPMC6103682.

37. Grootaert MOJ, Roth L, Schrijvers DM, De Meyer GRY, Martinet W. Defective Autophagy in Atherosclerosis: To Die or to Senesce? Oxid Med Cell Longev. 2018;2018:7687083. Epub 2018/04/24. doi: 10.1155/2018/7687083. PubMed PMID: 29682164; PubMed Central PMCID: PMCPMC5846382.

38. Torisu K, Singh KK, Torisu T, Lovren F, Liu J, Pan Y, et al. Intact endothelial autophagy is required to maintain vascular lipid homeostasis. Aging Cell. 2016;15(1):187–91. Epub 2016/01/20. doi: 10.1111/acel.12423. PubMed PMID: 26780888; PubMed Central PMCID: PMCPMC4717267.

39. Blazer AD, Clancy RM. ApoL1 and the Immune Response of Patients with Systemic Lupus Erythematosus. Curr Rheumatol Rep. 2017;19(3):13. Epub 2017/03/08. doi: 10.1007/s11926-017-0637-9. PubMed PMID: 28265848.

40. Thomson R, Finkelstein A. Human trypanolytic factor APOL1 forms pH-gated cation-selective channels in planar lipid bilayers: relevance to trypanosome lysis. Proc Natl Acad Sci U S A. 2015;112(9):2894–9. Epub 2015/03/03. doi: 10.1073/pnas.1421953112. PubMed PMID: 25730870; PubMed Central PMCID: PMCPMC4352821.

41. Kroemer G, Jaattela M. Lysosomes and autophagy in cell death control. Nat Rev Cancer. 2005;5(11):886–97. Epub 2005/10/22. doi: 10.1038/nrc1738. PubMed PMID: 16239905.

42. Cadwell K. Crosstalk between autophagy and inflammatory signalling pathways: balancing defence and homeostasis. Nat Rev Immunol. 2016;16(11):661–75. doi: 10.1038/nri.2016.100. PubMed PMID: 27694913; PubMed Central PMCID: PMCPMC5343289.

43. Ma L, Chou JW, Snipes JA, Bharadwaj MS, Craddock AL, Cheng D, et al. APOL1 Renal-Risk Variants Induce Mitochondrial Dysfunction. J Am Soc Nephrol. 2017;28(4):1093–105. doi: 10.1681/ASN.2016050567. PubMed PMID: 27821631; PubMed Central PMCID: PMCPMC5373457.

44. Lan X, Wen H, Saleem MA, Mikulak J, Malhotra A, Skorecki K, et al. Vascular smooth muscle cells contribute to APOL1-induced podocyte injury in HIV milieu. Experimental and molecular pathology. 2015;98(3):491–501. doi: 10.1016/j.yexmp.2015.03.020. PubMed PMID: 25796344.

45. Kupin WL. Viral-Associated GN: Hepatitis C and HIV. Clin J Am Soc Nephrol. 2017;12(8):1337–42. Epub 2016/11/01. doi: 10.2215/CJN.04320416. PubMed PMID: 27797895; PubMed Central PMCID: PMCPMC5544506.

